# Monomorphic *Trypanozoon*: towards reconciling phylogeny and pathologies

**DOI:** 10.1101/2021.04.14.439642

**Authors:** Guy Oldrieve, Mylène Verney, Kamil S. Jaron, Laurent Hébert, Keith Matthews

## Abstract

2.

*Trypanosoma brucei evansi* and *Trypanosoma brucei equiperdum* are animal infective trypanosomes conventionally classified by their clinical disease presentation, mode of transmission, host range, kDNA composition and geographic distribution. Unlike other members of the subgenus *Trypanozoon*, they are non-tsetse transmitted and predominantly morphologically uniform (monomorphic) in their mammalian host. Their classification as independent species or subspecies has been long debated and genomic studies have found that isolates within *T. b. evansi* and *T. b. equiperdum* have polyphyletic origins. Since current taxonomy does not fully acknowledge these polyphyletic relationships, we re-analysed publicly available genomic data to carefully define each clade of monomorphic trypanosome. This allowed us to identify, and account for, lineage specific variation. We included a recently published isolate, IVM-t1, which was originally isolated from the genital mucosa of a horse with dourine and typed as *T. equiperdum*. Our analyses corroborate previous studies in identifying at least four distinct monomorphic *T. brucei* clades. We also found clear lineage specific variation in the selection efficacy and heterozygosity of the monomorphic lineages, supporting their distinct evolutionary histories. The inferred evolutionary position of IVM-t1 suggests its reassignment to the *T. b. evansi* type B clade, challenging the relationship between the *Trypanozoon* species, the infected host, mode of transmission and the associated pathological phenotype. The analysis of IVM-t1 also provides the first evidence of the expansion of *T. b. evansi* type B, or a 5^th^ monomorphic lineage represented by IVM-t1, outside of Africa, with important possible implications for disease diagnosis.

**Impact statement:** *Trypanosoma brucei* are unicellular parasites typically transmitted by tsetse flies. Subspecies of *T. brucei* cause human African trypanosomiasis and the animal diseases, nagana, surra and dourine. *T. b. evansi* and *T. b. equiperdum* have branched from *T. brucei* and, by foregoing tsetse transmission, expanded their geographic range beyond the sub-Saharan tsetse belt. These species can only reproduce asexually and exhibit morphological uniformity in their host (‘monomorphism’). *T. b. evansi* and *T. b. equiperdum* have historically been classified based on fragmentary information on the parasites’ transmission routes, geographic distribution, kDNA composition and disease phenotypes. Our analysis of genome sequencing data from monomorphic *T. brucei* supports at least four independent origins with distinct evolutionary histories. One isolate, IVM-t1, typed as *T. equiperdum*, is a closer relative to *T. b. evansi*, highlighting the risk of using pathognomonic descriptors for subspecies assignment. We show clear lineage specific variation in the selection efficacy in monomorphic *T. brucei*. Using the evolutionary relationships between lineages, we suggest it would be beneficial to reconcile phylogeny and pathology in monomorphic trypanosomes.

**Data summary:** The data used in this study is available from the Sequence Read Archive or the Wellcome Sanger Institute. The accessions can be found in Supplementary file 1.

## 5. Introduction

The *Trypanozoon* subgenus (*Trypanosoma brucei* spp.) contains parasites of medical, veterinary and economic significance. In their mammalian form, developmentally competent (pleomorphic) trypanosomes transition from a proliferative ‘slender’ form to a cell-cycle arrested ‘stumpy’ form, adapted for survival in the midgut of its vector, the tsetse fly (*Glossina* spp.). Progression to the stumpy form occurs in a density dependent manner, mediated by a stumpy induction factor (1–4). Some *Trypanozoon* have a reduced ability to transition from the slender to stumpy morphotype and so are described as ‘monomorphic’. In the field, monomorphic *Trypanozoon* were historically classified as independent species, *T. equiperdum* and *T. evansi*, due to their distinct modes of transmission, geographic distribution, disease phenotype and host range (5). These monomorphic trypanosomes can infect livestock, and are currently implicated in causing dourine and surra, respectively (6).

More recently, it was proposed that *T. evansi* and *T. equiperdum* are subspecies of *T. brucei* (*T. b. evansi* and *T. b. equiperdum*) which had lost part or all of their kDNA, the parasites’ mitochondrial genome that encodes respiratory components required for viability in the tsetse fly vector (7). Whole genome comparisons found that monomorphism arose independently on at least four separate occasions, and further monomorphic isolates could be continuously emerging from pleomorphic *T. brucei* in the field. However, *T. b. evansi* and *T. b. equiperdum* are polyphyletic and can be assigned into at least four independently derived lineages, such that their subspecific names do not describe the evolutionary relationships between the different monomorphic *Trypanozoon* (8–10). *T. b. evansi* type A and *T. b. evansi* type B originate from *T. brucei* in Western and Central Africa whilst *T. b. equiperdum* type OVI and *T. b. equiperdum* type BoTat evolved from *T. brucei* in Eastern Africa (9). Whilst many naming conventions exist, for the remainder of this article we will use the proposition by Cuypers *et al*. (2017) (9), which currently most accurately describes the polyphyletic nature of monomorphic trypanosomes (i.e. *T. b. evansi* type A, *T. b. evansi* type B, *T. brucei, T. b. equiperdum* type OVI and *T. b. equiperdum* type BoTat).

The four *T. brucei* lineages converged on a monomorphic phenotype accompanied by a switch from cyclical to mechanical transmission. All of the monomorphic subspecies display a reduction or removal of their kDNA alongside an inability to complete their life cycle in their vector, locking these parasites into a tsetse-independent transmission mode (11,12). Current evidence suggests that *T. b. evansi* type A and *T. b. evansi* type B predominantly rely on transmission via biting flies (e.g., tabanids and *Stomoxys*), whilst *T. b. equiperdum* type OVI and *T. b. equiperdum* type BoTat are sexually transmitted between Equidae. Neither are cyclically transmitted via the tsetse vector (6) and, as sexual reproduction occurs in the tsetse salivary gland (13,14), monomorphic trypanosomes are obligately asexual and proliferate via mitosis (15). Escape from transmission by the tsetse fly, whose range is restricted to sub-Saharan Africa, has facilitated the expansion of monomorphic trypanosomes to other regions of Africa, Asia, Europe and the Americas, although they have subsequently been eradicated from North America and limited to local outbreaks in Europe (16).

The use of disease pathology, host species, geographic range and kDNA composition can complicate species classification where distinct lineages have converged on a phenotype. Research that treats polyphyletic lineages as a single group may miss more subtle, but important, differences between lineages. Here we have re-analysed existing genomic data from publicly available monomorphic *Trypanozoon* isolates to confirm their evolutionary relationships. This analysis included a recent isolate from Mongolia, IVM-t1, derived from the genital mucosa of a horse and classified as *T. equiperdum* based on its clinical disease symptoms and host species (17,18).

Through re-analysing publicly available whole genome data, we found at least four groups of monomorphic *T. brucei* with independent origins, consistent with previously published phylogenies (8,9). Our analysis concludes that currently IVM-t1 forms a clade with *T. b. evansi* type B, despite its clinical presentation being more typical of the conventional description of *T. equiperdum*. The presence of IVM-t1 in the genital mucosa of a horse with signs of dourine supports the hypothesis that there is considerable plasticity in the mode of transmission, host range and clinical presentation of monomorphic *Trypanozoon* strains of distinct origins, as was suggested by Brun *et al*. and Carnes *et al*. (8,19). These findings exemplify the need to classify monomorphic trypanosomes based on genetic information, unbiased by the mode of transmission, disease presentation, kDNA composition and host range. We also identified lineage specific variation in the heterozygosity and efficacy of selection of the four independent monomorphic lineages, highlighting the importance of characterising phylogeny-informed lineages. Finally, we note that the ancestor of the *T. b. evansi* type B clade, or a 5^th^ monomorphic lineage represented by IVM-t1, extended its geographic range outside of Africa.

## 6.1 Methods

### Variant calling

Publicly available genome data was accessed from the Sequence Read Archive (SRA) (20) and the Wellcome Sanger Institute. Samples sequenced with older technologies, such as solid-state ABI, were excluded. When PacBio and Illumina data was available for the same sample, Illumina data was used preferentially to standardise the comparison.

The *T. brucei* EATRO 1125 Antat 1.1 90:13 (21) genome was sequenced as part of this study. DNA was extracted using the DNeasy Blood & Tissue Kit with an RNAse A step (Qiagen) following the manufacturer’s instructions. The DNA was sequenced (HiSeq 4000) and cleaned by BGI, Hong Kong (7,723,274 reads at 150 base pair length). The *T. brucei* EATRO 1125 Antat 1.1 90:13 raw data has been submitted to SRA (PRJNA720808). The complete list of genomes analysed in this study, including their accession IDs, are summarised in the supplementary file 1.

The quality of the raw reads were analysed with fastqc (v:0.11.9) and subjected to quality trimming with trimmomatic (v0.39) (22). The reads were trimmed with the following filters: SLIDINGWINDOW:4:20, ILLUMINACLIP:adapters.fa:2:40:15, MINLEN:25. The trimmed reads were aligned to the *T. brucei* TREU927/4 V5 reference genome (23) with bwa-mem (v:0.7.17-r1188) (24). The reads were prepared for variant calling by following the GATK4 (v:4.1.4.1) best practices pipeline which included marking duplicate reads (25,26). The read recalibration step was performed by initially calling variants on un-calibrated reads with GATK4’s haplotype caller (27). The top 20% highest confidence calls from the first round were used as the confident call set to re-calibrate the raw bam files. Variants were then re-called on the recalibrated bam files with GATK4’s haplotype caller (27).

The variants were combined and filtered with the following stringent cut-offs, in keeping with GATK’s best practices pipeline and previous studies (9). SNPs were filtered by quality by depth (< 2.0), quality score (< 500.0), depth (<5.0), strand odds ratio (> 3.0), Fisher’s exact (> 60.0), mapping quality (< 40.0), mapping quality rank sum (< -12.5), read position rank sum (< -8.0), window size (10) and cluster size (3). Indels were filtered on their quality by depth (< 2.0), quality (< 500.0), Fisher’s exact (> 200.0) and read position rank sum (< - 20.0).

### Phylogenetic analysis

The filtered variants, described above, were filtered again to retain sites where a genotype had been called in every sample, using VCFtools (v:0.1.16) (28). These sites were analysed in two ways. The first was based on SNPs which occurred across all of the sites in the *T. brucei* TREU927/4 V5 reference genome and the second from SNPs which occurred in a CDS of a gene (excluding pseudogenes) found on one of the 11 megabase chromosomes (23). For both of these analyses, a concatenated alignment of each variant was extracted using VCF-kit (v:0.1.6) (29). IQ-TREE (v:2.0.3) (30) was used to create a maximum likelihood tree from homozygous variant sites. Within the IQ-TREE analysis, a best-fit substitution model was chosen by ModelFinder using models which included ascertainment bias correction (MFP+ASC) (31). ModelFinder identified TVM+F+ASC+R2 as the best fit for both of the alignments which were subjected to 1000 ultrafast bootstraps generated by UFBoot2 (32). The consensus trees were visualised and annotated using iTOL (v:5) (33).

### Genome content

Raw reads were used to predict the heterozygosity, GC content and genome size of each isolate. This analysis was performed using a k-mer counting based approach with Jellyfish (v:2.3.0) (34) and Genomescope (v:1). A k-mer size of 21 was used (35).

Non-synonymous and synonymous SNP ratios (dN/dS) can be used to estimate the selection pressure upon an organism. The filtered SNPs identified for the phylogenetic analysis was split by isolate using bcftools (v:1.9) (36) and these were then filtered to remove all non-variant sites using GATK4 SelectVariants (v:4.1.9.0). The reverse complement of each variant call format (VCF) file was generated using SNPGenie (v: 2019.10.31) (37). SNPGenie within-pool analysis was then completed on each isolate for SNPs found in the longest CDS site of each gene, excluding pseudogenes, on the 11 megabase chromosomes of the *T. brucei* TREU927/4 V5 reference genome (23). The full dN/dS results are available in Supplementary file 2.

### Molecular markers

Molecular marker sequences were downloaded from NCBI: SRA (Z37159.2), RoTat1.2 (AF317914), *evansi* VSG JN2118HU (AJ870487), cytochrome oxidase subunit 1 (CO1) (M94286.1:10712-12445) and NADH dehydrogenase subunit 4 (NADH4) (M94286.1:12780-14090). bwa-mem (v:0.7.17-r1188) (24) was used to align raw reads to the molecular markers. A minimum overlap of 50 base pairs between the read and target sequence was used. The molecular marker presence was confirmed by counting the number of bases in the target sequence covered by reads, representing the breadth of coverage, with samtools mpileup (38). Orphan reads were counted and read pair overlap detection was disabled. The breadth of coverage percentage was visualised with pheatmap (v:1.0.12) (39). The ATP synthase γ subunit (Tb927.10.180) was screened for lineage specific variants identified in the variant calling step.

Unless stated otherwise, all figures were plotted using ggplot2 (v:3.3.0) (40) and ggrepel (v:0.8.1) (41) in R (v:3.6.1) (42).

## 6.2 Results

### Monomorphism arose independently at least four times

To determine the evolutionary relationships between monomorphic *Trypanozoon*, genomic data from 17 isolates were aligned to the *T. brucei* TREU 927/4 reference genome. This included publicly available data from monomorphic (n=9) and pleomorphic isolates (n=8). Across the *T. brucei* TREU 927/4 v5 reference genome, 472,794 variant sites (574,775 unique variant alleles, 370,154 SNPs and 204,621 indels) passed the strict quality filtering steps. These sites were filtered further to identify 244,013 homozygous variant SNPs across the genome. 91,853 of these SNPs were present in a CDS on one of the 11 megabase chromosomes of the *T. brucei* TREU 927/4 v5 reference genome. The SNPs across the whole genome and those in a CDS were used to generate two unrooted phylogenetic trees.

Monomorphic *T. brucei* form at least four independent clades (Fig. 1). Our results corroborate previous findings (8,9) which identified that the *T. b. equiperdum* type OVI clade arose in Eastern Africa and displays minimal variation between the isolates. *T. b. equiperdum* type BoTat is separated from other *T. b. equiperdum* isolates and represents an Eastern African isolate of distinct origin. Trypanosomes designated *T. b. evansi* also form two discrete clades of Western and Central African origin, with STIB805, RoTat1.2 and MU09 (*T. b. evansi* type A) displaying low genetic diversity and being distinct from MU10 (*T. b. evansi* type B).

**Figure 1:**
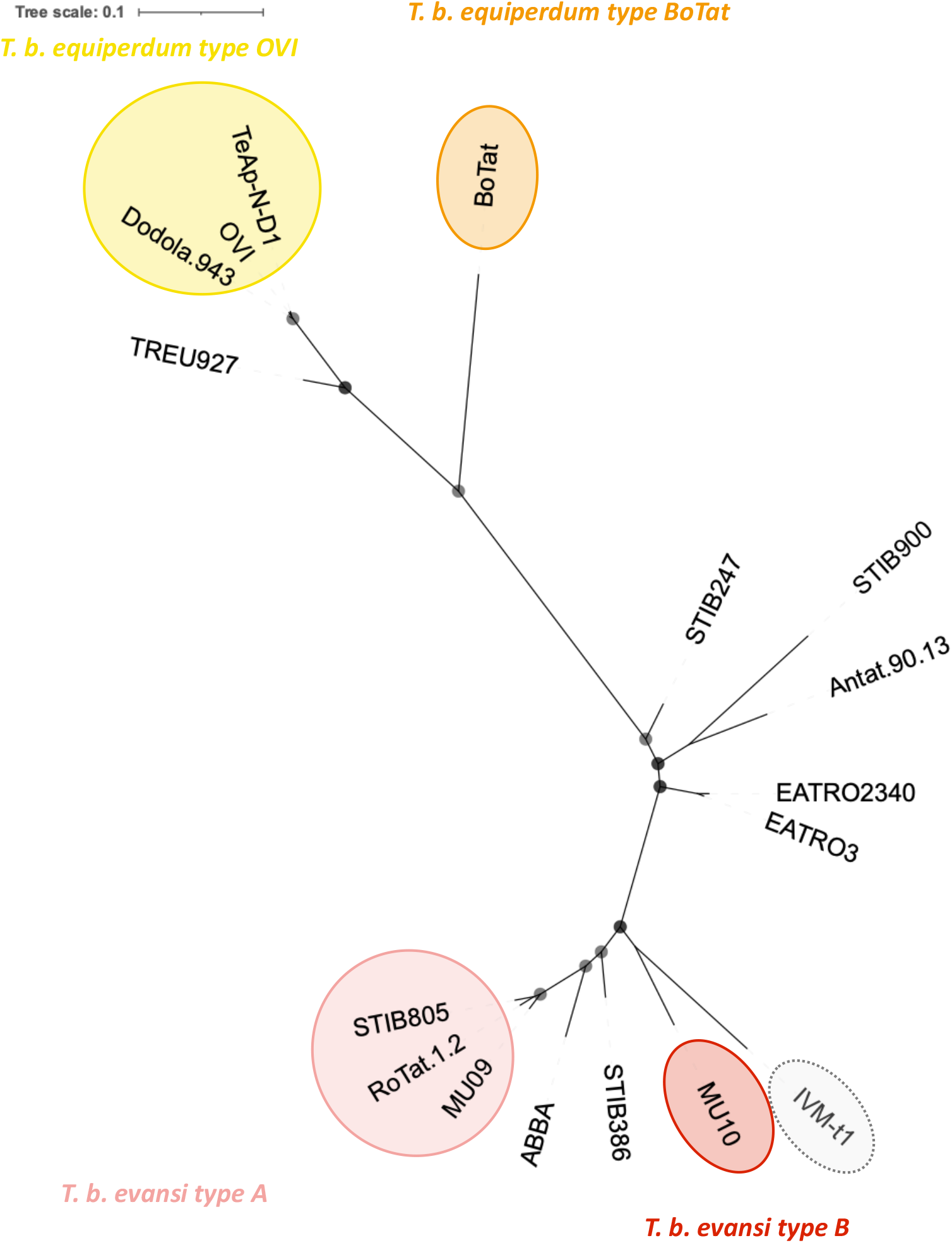
An unrooted phylogenetic tree created with 244,013 homozygous variant SNPs found across the *T. brucei* TREU 927/4 reference genome. The tree was built using a TVM+F+ASC+R2 model. Bootstrap confidence is reported by the size of grey circles. All bootstrap values were 100 and so each circle is the same size. Monomorphic genomes form four distinct lineages which have expanded from Eastern (*T. b. equiperdum* type OVI and *T. b. equiperdum* type BoTat) and Western/Central Africa (*T. b. evansi* type A and *T. b. evansi* type B) (9). IVM-t1 was originally typed as *T. b. equiperdum* but groups here with *T. b. evansi* type B.

Interestingly, the Mongolian isolate IVM-t1, with infection and disease characteristics similar to *T. b. equiperdum*, branched with MU10 which is of West African origin. This contrasts with its previous designation as a *T. equiperdum* isolate, whose ancestors originated from Eastern Africa (9) (Fig. 1). Nonetheless, whilst IVM-t1 and MU10 shared a more recent common ancestor with each other than with any other strain in this analysis, MU10 and IVM-t1 have diverged considerably. Should pleomorphic *T. brucei* isolates be identified which divide the clade composed of MU10 and IVM-t1, the isolates would represent independent clades.

The results generated from SNPs found across the whole genome were similar to a tree built from SNPs found in only the CDS, with the former displaying a slight reduction in bootstrap confidence (Fig. S1).

### There is quantifiable variation in the genetic diversity, and efficacy of selection between the four asexual monomorphic clades

Asexuality is expected to reduce the heterozygosity of the lineage, although this can be lineage specific (43). Asexuality can also reduce the efficacy with which selection can act, as reviewed by Otto, 2021 (44). Efficacy of selection can be estimated by calculating the ratio of nonsynonymous to synonymous variants (dN/dS) compared to a reference genome.

Our analysis demonstrated that monomorphic genomes generally have a lower heterozygosity compared to the genomes derived from pleomorphic isolates (Fig. 2 & Fig.S2). Notable outliers in this analysis includes MU10 (*T. b. evansi* type B) which has a heterozygosity value closer to that of pleomorphic genomes. The higher dN/dS ratio observed in *T. b. equiperdum* type OVI is in contrast to the other monomorphic lineages, highlighting the lineage specific variation, and evolutionary histories, of monomorphic *Trypanozoon*.

**Figure 2:**
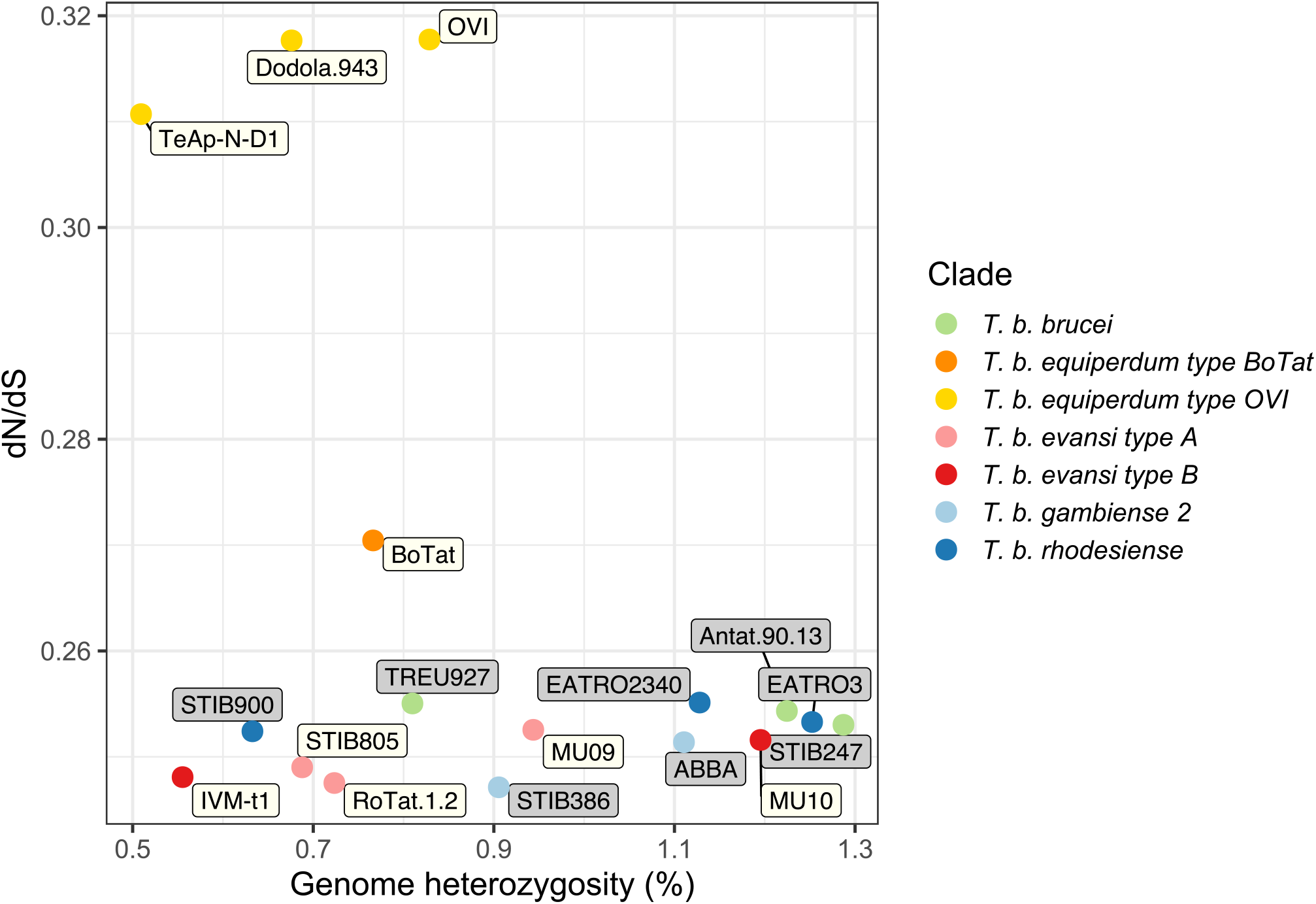
Whole genome heterozygosity and dN/dS ratio of SNPs present in the longest CDS of every annotated gene, excluding pseudogenes, found on one of the 11 megabase chromosomes of the *T. brucei* TREU 927/4 reference genome. The values were calculated for all publicly available monomorphic isolates and representative pleomorphic isolates. Each point is coloured by clade and the label colour represents a pleomorphic (grey) or monomorphic (white) isolate.

Interestingly, IVM-t1 has the second lowest heterozygosity and one of the lowest dN/dS ratios in this analysis, more similar to that of *T. b. evansi* type A and *T. b. evansi* type B and in contrast to the levels observed in *T. b. equiperdum* type OVI and *T. b. equiperdum* type BoTat (Fig. 2). Raw reads from *T. b. brucei* TREU 927/4 were included in the analysis. The variants called against its own genome assembly represent heterozygous loci and misaligned reads. *T. b. equiperdum* BoTat has the longest branch length in this analysis but has a far lower dN/dS ratio than the *T. b. equiperdum* type OVI clade (Fig. S3). The low dN/dS of *T. b. brucei* TREU 927/4 called against its own genome and the pattern of branch length and dN/dS ratio highlights that high dN/dS of *T. b. equiperdum* type OVI is not the result of the evolutionary distance between the clade and the reference genome.

### Genetic markers corroborate phylogenetic and genome content analysis

The occurrence of established molecular markers for different *Trypanosoma brucei* subspecies was compared between the isolates to further validate whole genome analysis. The markers’ occurrence in each isolate was visualised in a hierarchical clustered heatmap. The clustering highlights a clear distinction between the monomorphic lineages and furthermore, with just five molecular markers, it is possible to recreate a similar pattern to the phylogenetic tree which is based on whole genome sequencing data (Fig. 1 & 3).

As our phylogenetic and genome content analysis identified a potential discrepancy between the disease description of IVM-t1 and its genotype, particular interest was paid to this isolate. Firstly, being akinetoplastic at the point of sequencing, IVM-t1 lacks coverage of the mitochondrial maxicircle genes cytochrome oxidase subunit 1 (CO1) (M94286.1:10712-12445) and NADH dehydrogenase subunit 4 (NADH4), as do *T. b. evansi* type A and *T. b. evansi* type B. However, when IVM-t1 was initially isolated, a PCR for NADH4 found that it contained the gene, albeit with a faint signal (17). Therefore, maxicircle presence in IVM-t1 appears to have been unstable, as is the case in kDNA independent isolates. IVM-t1 reportedly became akinetoplastic after long term culture adaptation, as has been observed in other monomorphic isolates (45,46). In contrast to IVM-t1, CO1 and NADH4 are present in *T. b. equiperdum* type OVI and *T. b. equiperdum* type BoTat genomes.

Secondly, IVM-t1 lacks the RoTat 1.2 VSG (AF317914.1), which is diagnostic for *T. b. evansi* type A (47), but does have sequence coverage of VSG JN2118HU (AJ870487.1) which is present in *T. b. evansi* type B, along with some *T. b. brucei* strains (48,49). In contrast, JN2118HU is absent in *T. b. evansi* type A, *T. b. equiperdum* type OVI *and T. b. equiperdum* type BoTat (Fig. 3).

**Figure 3:**
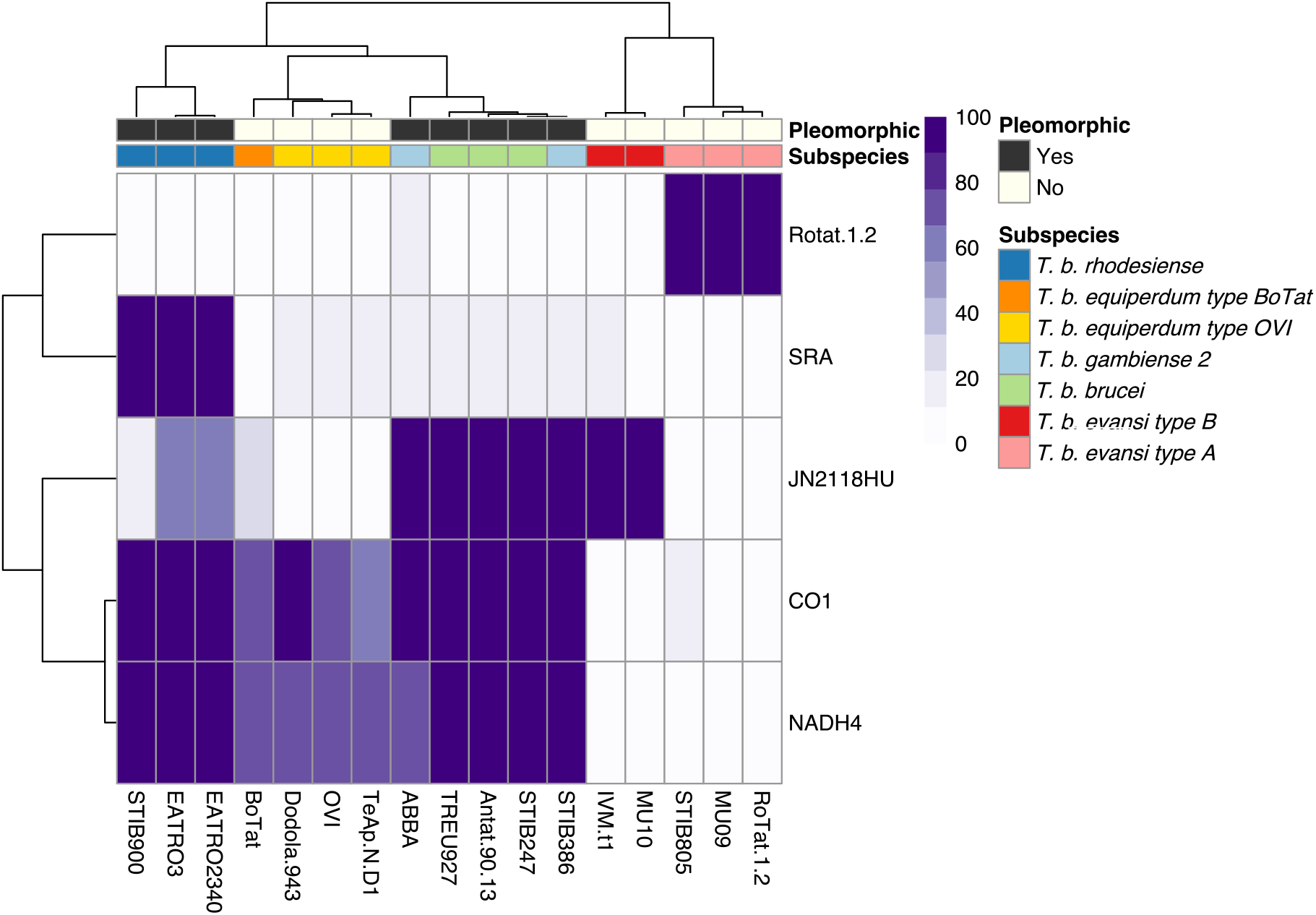
Occurrence of individual genetic markers corroborate phylogenetic and whole genome analysis which highlights at least four independent monomorphic lineages. Genetic markers: SRA (Z37159.2), RoTat1.2 VSG (AF317914.1), JN2118HU VSG (AJ870487.1), cytochrome oxidase subunit 1 (M94286.1:10712-12445) and NADH4 (M94286.1:12780-14090). The scale represents the percentage of the marker covered by sequencing reads.

Thirdly, IVM-t1 does not have the M282L ATP synthase γ subunit mutation which has been characterised in other *T. b. evansi* type B genomes, such as MU10 (Supplementary file 3) (49,50). This mutation is unable to compensate for kDNA loss (50). However, IVM-t1 does have two homozygous mutations within that gene which are absent from all other isolates in this analysis (C-terminal genomic codons G817T and A898G) although both are synonymous and so predicted not to influence protein function.

In combination, these analyses further support the separation of IVM-t1 from the *T. b. equiperdum* type OVI and *T. b. equiperdum* type BoTat lineages and also highlights differences to *T. b evansi* MU10 lineage despite the phylogenetic relationship between the isolates.

## 6.3 Discussion

*Trypanozoon* phylogeny has historically been based on clinical disease pathology, mode of transmission, geographic range, host species range and kDNA composition, which has complicated the classification of monomorphic trypanosomes and fuelled a long-standing debate in the literature (5). Here we re-analyse the molecular phylogeny of monomorphic trypanosomes, providing support for the complete separation of these isolates into at least four clades based on their evolutionary relationships. Our results are consistent with previously published phylogenies of monomorphic *T. brucei* subspecies (Fig.1) (8,9). We consider it is important to classify the monomorphic lineages through their evolutionary relationships because, although they have converged upon a monomorphic phenotype, lineage specific variation could be missed if their polyphyletic origin is not fully acknowledged.

Variation in heterozygosity can be associated with a transition to asexuality. Asexual taxa can present high heterozygosity if the lineage arose from a hybrid origin, but other types of origin usually lead to a reduction in heterozygosity (43). Monomorphic strains have lost their tsetse transmission ability. Given that meiotic events occur in tsetse salivary glands, monomorphic strains are obligately asexual and proliferate via mitosis (13–15). This asexuality is apparent in the reduction of heterozygosity observed in the majority of monomorphic isolates (Fig. 2 & Fig. S2). Such reduction in heterozygosity can occur via mitotic recombination and/or gene conversion, with gene conversion having been observed in another asexual *T. brucei* subspecies, *T. b. gambiense* group 1 (51). Gene conversion is proposed to reduce or even completely stop the mutational attrition associated with asexuality, and it can occur at a faster rate than the accumulation of spontaneous mutations (52) (Fig.2 & Supplementary Fig.2).

Notably, *T. b. equiperdum* type OVI and, to an extent, *T. b. equiperdum* type BoTat isolates have low heterozygosity and a higher dN/dS ratio than other clades, indicative of a smaller efficacy of purifying selection in removing deleterious alleles. In contrast, the efficacy of selection in *T. b. evansi* type A and *T. b. evansi* type B are closer to that observed in pleomorphic lineages. Further, IVM-t1, originally typed as *T. equiperdum*, has one of the lowest dN/dS ratios, more indicative of *T. b. evansi* type A or *T. b. evansi* type B. The efficacy of selection can be influenced by the effective population size which is linked to events during the evolutionary history of a lineage such as population bottlenecks and variation in the mode of inheritance (53). Asexuality is predicted to reduce the efficacy with which selection can act (44). Hence, the observed lineage specific variation between selection efficacy could be associated with a different length of time as an asexual lineage as the predicted build-up of deleterious mutations is a gradual process, if not completely counteracted by processes like gene conversion. Overall, the analyses of heterozygosity and dN/dS ratio support the different monomorphic lineages displaying contrasting evolutionary histories.

The extremely low heterozygosity of IVM-t1 highlights that when the sample was sequenced, it did not comprise a mixed infection. Furthermore, its low heterozygosity does not support a hybridisation-based event at the emergence of the IVM-t1 lineage (43). Therefore, the long branch and discrepancy in heterozygosity between IVM-t1 and MU10 could be due to an expansion in diversity of the MU10 branch or independent origins of monomorphism within the MU10/ IVM-t1 clade. This divergence could have facilitated the distinct transmission mechanism and host range displayed by IVM-t1 with respect to *T. b. evansi* type B such as MU10. However, at present IVM-t1 and MU10 group as a separate clade and share the monomorphic phenotype. As such IVM-t1 is currently most accurately described as *T. b. evansi* type B. As more *T. brucei* are isolated and sequenced, it may be more accurate to define IVM-t1 as a separate clade. In this case, IVM-t1 would represent a 5^th^ independent emergence of monomorphism in *T. brucei*.

The potential expansion of monomorphic lineages, along with the isolation of IVM-t1 from the genital mucosa of a horse with signs of dourine suggest it could be beneficial to reconcile phylogeny and disease (17). To fully uncouple the link between phylogeny and disease, studies will be required on the direct mode of transmission of these isolates. For instance, although the presence of IVM-t1 in the genital mucosa of a horse with signs of dourine points towards sexual transmission, it cannot be ruled out that the initial infection was a coinfection of IVM-t1 with an independent *T. b. equiperdum* type OVI or *T. b. equiperdum* type BoTat isolate which caused the dourine symptoms but was not recovered after culture. If plasticity in the mode of transmission is established, dourine and surra would best refer to the disease presentation and not the causative agent, particularly where limited clinical information is available for an isolate, precluding an understanding of any variable disease manifestation between individual animals (6).

The emergence of at least four independent monomorphic lineages suggests there could be a selective advantage to monomorphism, at least in the short term. Since monomorphic lineages lose the growth control inherent in the generation of stumpy forms, they display an increase in parasitaemia which improves the chance of non-tsetse transmission when tsetse vectorial capacity is reduced. This adaption to a loss of cyclical transmission could also be the only option for those *Trypanozoon* isolates which become isolated from the tsetse fly through environmental change or geographic relocation of the host (7,54,55). Regardless of the potential adaptive advantage to monomorphism, monomorphic lineages could be constantly emerging from pleomorphic *T. brucei* populations across Africa that remain unidentified due to a lack of sampling. Indeed, as climate change rapidly alters the tsetse flies’ range (56), creating a potential selective advantage for mechanical transmission in areas where tsetse flies are no longer found, the rate of monomorphic *T. brucei* subspecies emergence could increase. Since monomorphic lineages have lost or reduced their growth control mechanism, they can be highly virulent, posing a threat to livestock in their country of origin and with the risk of escape outside traditional disease boundaries.

The analysis of IVM-t1 provides the first evidence that *T. b. evansi* type B, or a 5^th^ monomorphic lineage, has expanded its geographic range outside of Africa. Previously *T. b. evansi* type B has only been found in Kenya and Ethiopia (48,49,57). The presence of IVM-t1 outside of Africa could complicate the existing screening of animals exhibiting signs of dourine or surra. Currently, the molecular diagnosis of surra and dourine remains limited by the parasitaemia in infected hosts, which can be below the detection limit of parasitological tests and can even be below the detection limit of DNA tests, especially in Equidae and African cattle. Therefore, serological methods are prescribed by the World Organisation for Animal Health for surra and dourine diagnosis. In some regions and for some hosts, for which *T. b. evansi* type A strains are widely present, the use of serological tests based on the recognition of the specific RoTat 1.2 VSG, can provide good sensitivity and specificity (47). However, this gene is absent in *T. b. evansi* type B (Fig.3). Alternatively, surveillance of kDNA integrity remains a useful method of identification, which often aligns with the disease presentation despite these phenomena not being biologically linked. It is important to remember, however, that kDNA integrity comes with the risk of the independent appearance of dyskinetoplasty in multiple lineages or the spontaneous or progressive dyskinetoplasty observed after maintenance *in vitro* culture or in isolates indistinguishable at the level of the nuclear genome.

To avoid these complications, it is possible to rely on markers that are generic for all trypanosome subspecies. Indeed, given that treatment success depends mainly on the stage of the disease rather than the specific *Trypanozoon* lineage (6), we suggest that currently it remains preferable not to aim for a distinction between taxa within the *Trypanozoon* subgenus for first line diagnosis. However, genome sequencing is rapidly reducing in cost whilst improving in portability. Therefore, adaptive genome sequencing represents a promising method for screening of animals infected with monomorphic *Trypanozoon* (58).

We note that although the use of four clades (*T. b. evansi* type A, *T. b. evansi* type B, *T. b. equiperdum* type OVI and *T. b. equiperdum* type BoTat (9)) attempts to acknowledge the polyphyletic origin of monomorphic *Trypanozoon*, the use of *evansi* or *equiperdum* at the subspecific level is not based on the genetic relationship of the strains. This is the case even when phylotypes are utilised after *evansi* or *equiperdum*. Therefore, we suggest it would be beneficial to break the links between the fragmentary information available for taxonomy, disease phenotype, host range, mode of transmission and the extent of dyskinetoplasty in monomorphic *Trypanozoon*.

Instead, to fully acknowledge their polyphyletic origin and distinct evolutionary histories, we suggest the taxonomy of monomorphic *Trypanozoon* should be based on whole genome analysis alone, which is quantitative, non-subjective and can be assessed from samples lacking detailed case records. A taxonomy based on the evolutionary relationships between isolates will assist future research by identifying lineage specific variation in monomorphic *Trypanozoon*. For example, we consider IVM-t1 is currently more accurately classified as a branch of *T. b. evansi* type B, rather than *T. equiperdum*. Similarly, isolates such as STIB818 (isolated China in 1979) and ATCC30023 (isolated in France in 1903) were initially classified as *T. equiperdum* but cluster with *T. b. evansi* (8,59). The disease manifestation and tissue specificity of IVM-t1 also suggests that *T. b. evansi* type B, or a 5^th^ monomorphic lineage, can manifest dourine symptoms with sexual transmission, although direct transmission evidence is needed to confirm this.

The incongruity between the parasites’ evolutionary position and the induced pathology, mode of transmission and tissue tropism highlights the potential for the host physiology and immune response to contribute to clinical disease manifestation, rather than being solely parasite driven (6). This provides an important exemplar of the potential distinction between the taxonomic position of monomorphic trypanosomes and the diseases they cause.

## 7. Author statements

### 7.1 Authors and contributions

Guy Oldrieve: 0000-0003-1428-0608

Mylène Verney: 0000-0002-0708-0045

Kamil Jaron: 0000-0003-1470-5450

Laurent Hébert: 0000-0002-5738-9913

Keith Matthews: 0000-0003-0309-9184

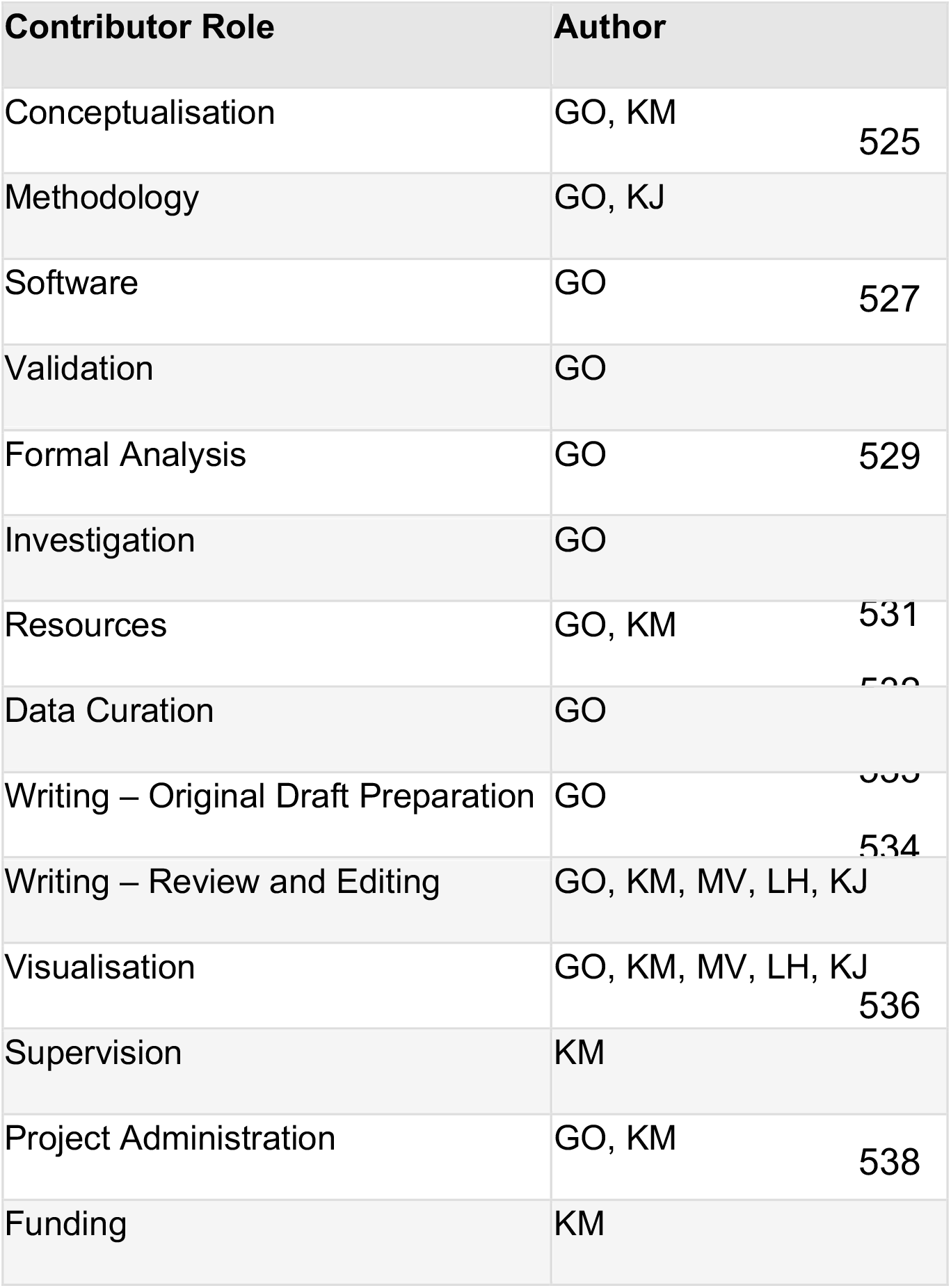

## 7.4 Ethical approval

Not applicable.

## 7.5 Consent for publication

Not applicable.

## 7.6 Acknowledgments

We thank Achim Schnaufer for his valuable insights and comments on the manuscript.

---

**Supplementary figure 1:**
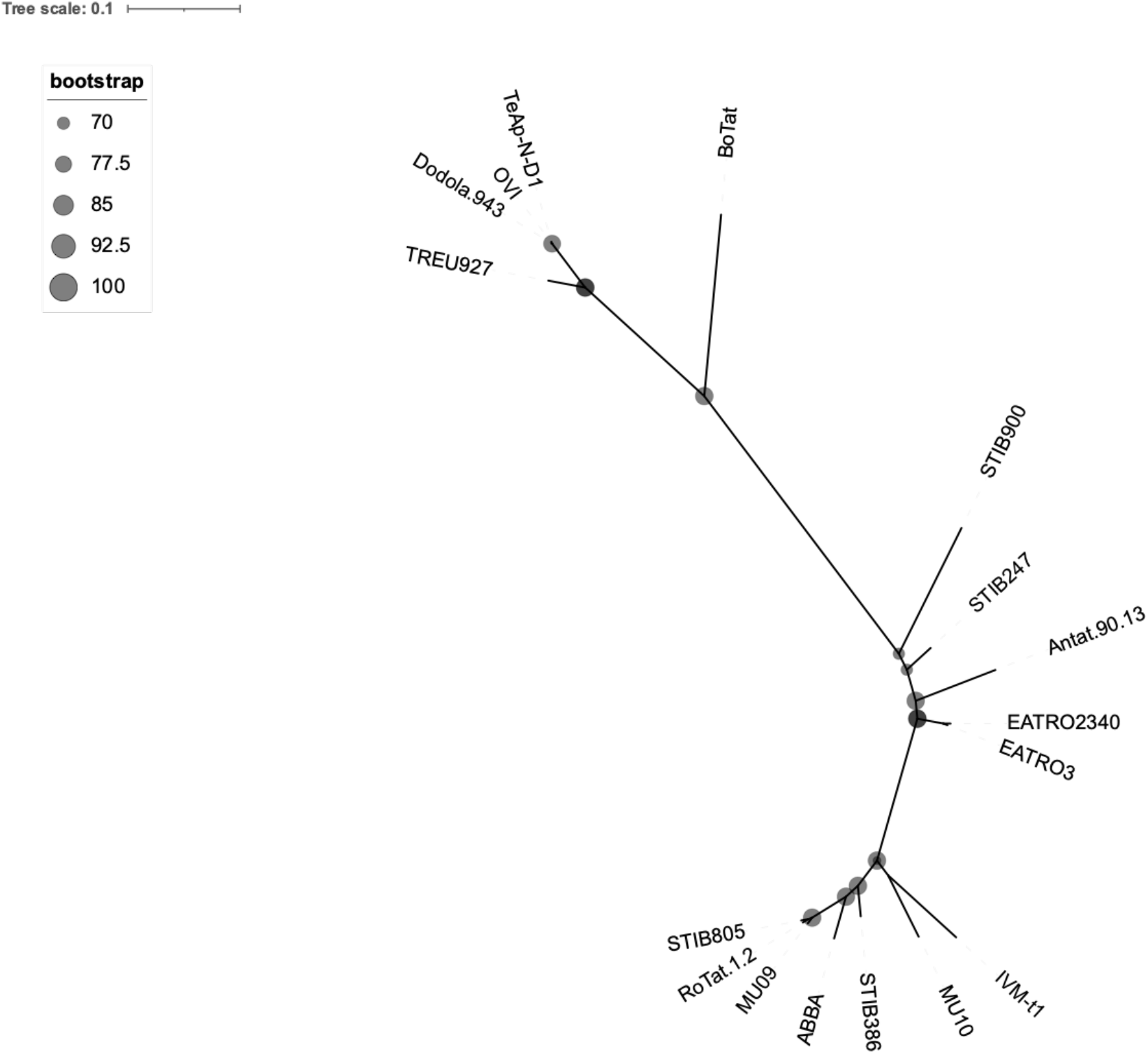
An unrooted phylogenetic tree created with 91,853 homozygous variant SNPs identified in a CDS on one of the 11 megabase chromosome of the *T. brucei* TREU 927/4 reference genome. The tree was built using the TVM+F+ASC+R2 model. Bootstrap values are reported by circle size. Monomorphic genomes form four distinct lineages which have expanded from Eastern (*T. b. equiperdum* type OVI and *T. b. equiperdum* type BoTat) and Western/Central Africa (*T. b. evansi* type A and *T. b. evansi* type B).

**Supplementary figure 2:**
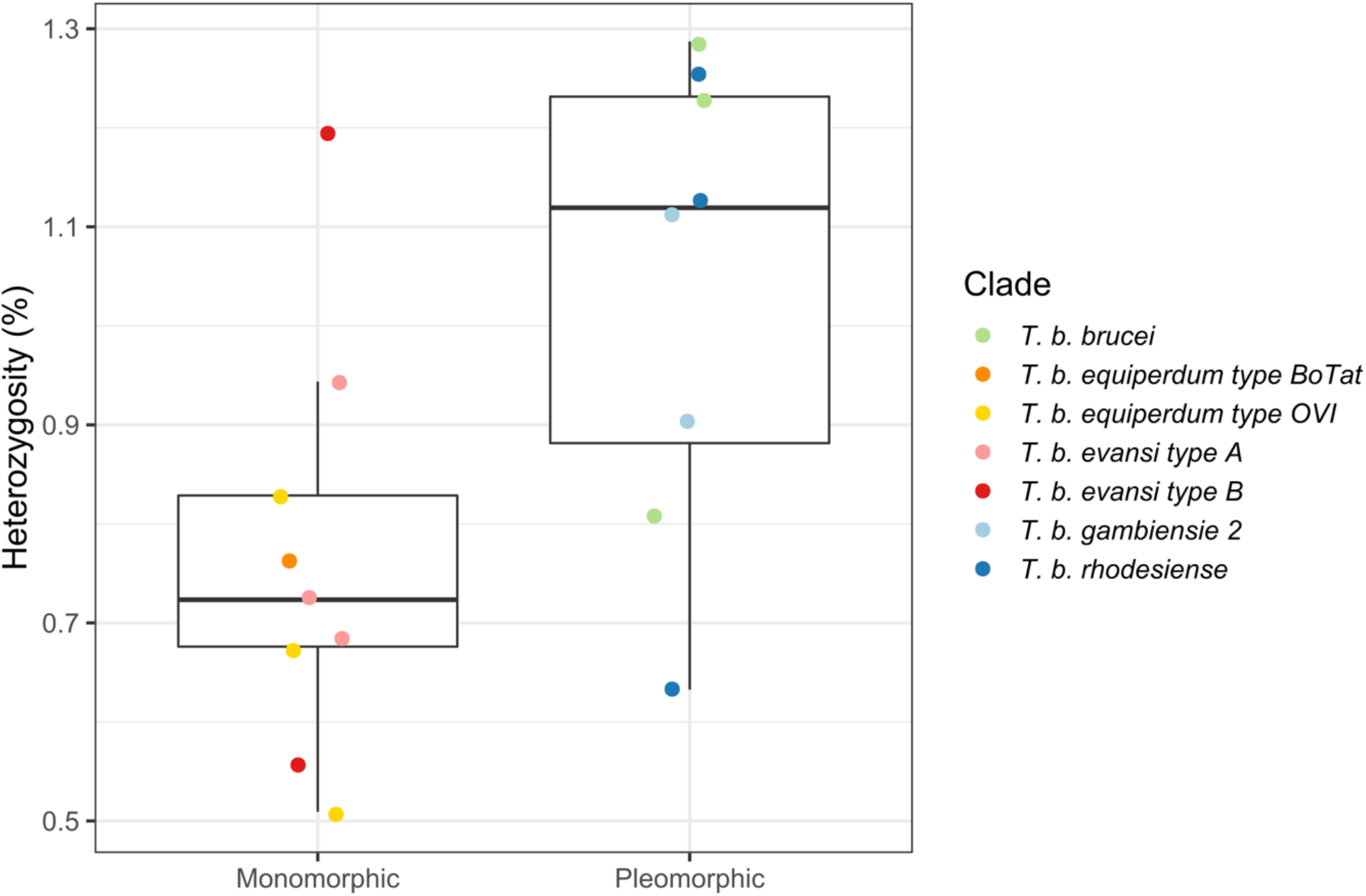
Whole genome heterozygosity of monomorphic and pleomorphic isolates.

**Supplementary figure 3:**
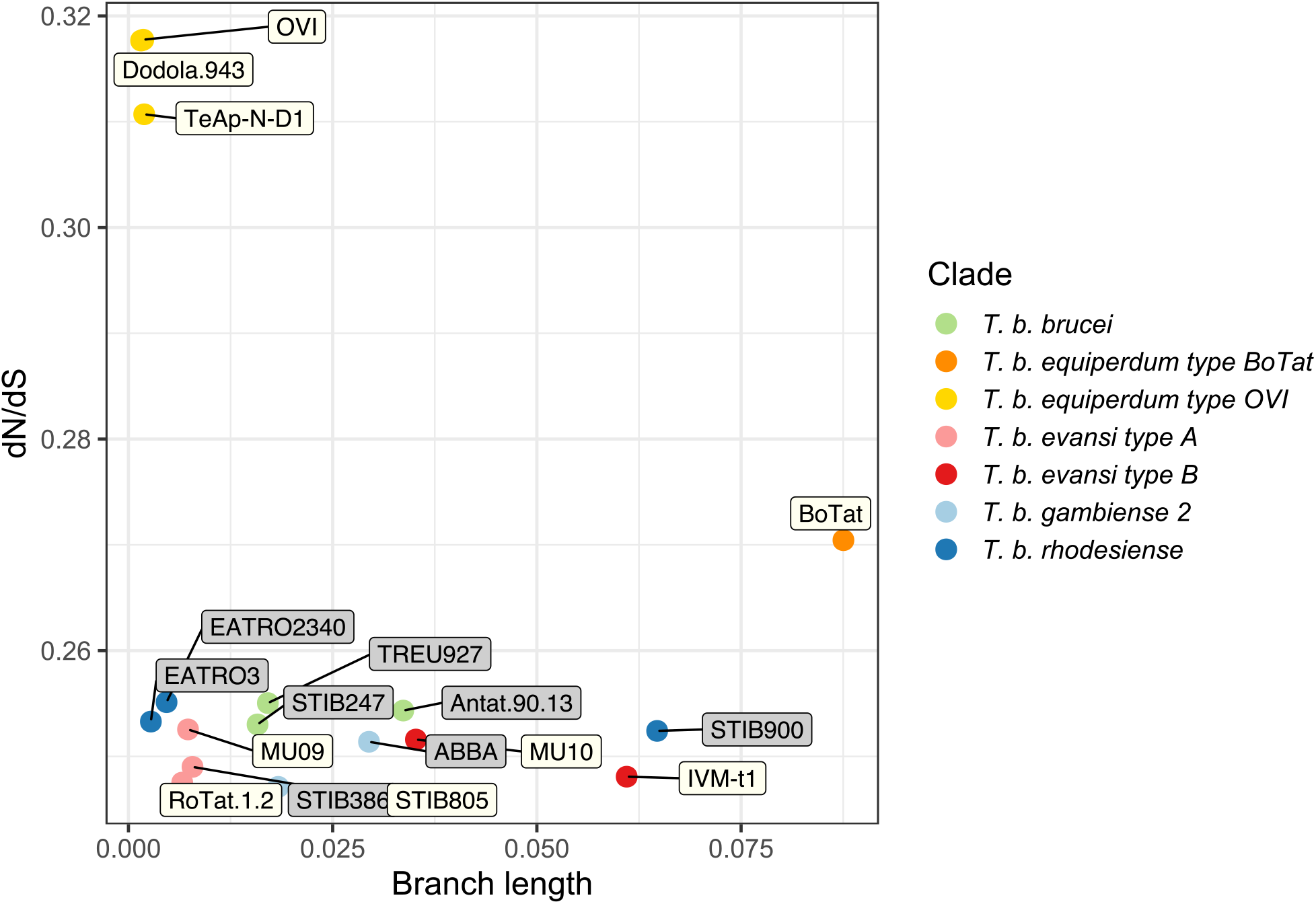
Branch length and dN/dS ratio of SNPs present in the CDS of genes found on the 11 megabase chromosomes of the *T. brucei* TREU 927/4 reference genome. The values were calculated for all publicly available monomorphic isolates and representative pleomorphic isolates. Each point is coloured by clade and the label colour represents a pleomorphic (grey) or monomorphic (white) isolate.

